# Diversity of sex chromosomes in Sulawesian medaka fishes

**DOI:** 10.1101/2022.02.28.482394

**Authors:** Satoshi Ansai, Javier Montenegro, Kawilarang W.A. Masengi, Atsushi J. Nagano, Kazunori Yamahira, Jun Kitano

**Author notes:** **Correspondence**, Satoshi Ansai, Graduate School of Life Sciences, Tohoku University, Japan.

## Abstract

Recent genetic and genomic studies have revealed tremendous diversity in sex chromosomes across diverse taxa. Although theoretical studies predict that sex chromosome evolution can drive the evolution of sexual dimorphism, empirical studies are still limited. A taxonomic group that shows diversity in both sex chromosomes and sexual dimorphism provides excellent opportunity to investigate the link between the evolution of sex chromosomes and sexual dimorphism. The medaka fishes (family Adrianichthyidae) exhibit both rapid sex chromosome turnovers and diversification of sexually dimorphic traits. In the present study, we investigated the sex chromosomes of 13 *Oryzias* species from Sulawesi, Indonesia, that have diversified in sexual dichromatism. Using pooled sequencing we found sex chromosomes in 9 species that all had XY systems, with a few species being possibly modified by multiple loci. Seven species (*O. woworae, O. asinua, O. wolasi, O. matanensis, O. celebensis, O. hadiatyae*, and *O. dopingdopingensis*) share linkage group (LG) 24 as sex chromosomes; however, they differed in the length and magnitude of sequence divergence between the X and Y chromosomes. The sex chromosome of *O. eversi* was LG4, which has not been reported as a sex chromosome in any other medaka species. In *O. sarasinorum*, LG16 and LG22 are associated with sex. Although LG16 was found to be sex-linked in another medaka species previously examined, the sex-determining regions did not overlap. Frequent turnovers and the great diversity of the sex chromosomes make Sulawesian medaka species a model system for investigating the roles of sex chromosome evolution in the diversification of sexual dimorphism.

## 1 INTRODUCTION

Recent genetic and genomic studies have revealed tremendous diversity in sex chromosomes across diverse taxa (Bachtrog et al., 2014). In several taxa, closely related species or even populations use different chromosomes for sex determination (Gammerdinger and Kocher, 2018; Miura, 2017; Myosho et al., 2015; Ross et al., 2009). Such sex chromosome turnover may be associated with the evolution of sexual dimorphism (Kitano and Peichel, 2012; Mank, 2009; Rice, 1984). A sexually antagonistic allele at an autosomal locus can drive the transposition of a pre-existing sex determining gene or the evolution of a new sex determining gene near that locus because it can resolve the intra-locus sexual conflict by enabling males and females to have different genotypes (Charlesworth and Charlesworth, 1980; van Doorn and Kirkpatrick, 2007). Although a few examples demonstrate that genes responsible for sexual dimorphism are located on newly evolved sex chromosomes (Kitano et al., 2009; Roberts et al., 2009), this hypothesis remains to be rigorously tested.

Alternatively, genes responsible for sexual dimorphism may evolve after the evolution of sex chromosomes. Recombination suppression is a universal feature of sex chromosomes, although the drivers of recombination suppression remain controversial (Beukeboom and Perrin, 2014; Jeffries et al., 2021; Lenormand and Roze, 2022; Ponnikas et al., 2018). Once sex chromosomes stop recombination, X and Y (or Z and W) chromosomes start to diverge in regulatory sequences, amino acid sequences, and/or the composition of genes (Lenormand and Roze, 2022; Yoshida et al., 2014; Zhou and Bachtrog, 2012). These differences may lead to the evolution of sexual dimorphism. Particularly, mutations that can resolve intra-locus sexual conflict may tend to accumulate in such non-recombining regions (Rice, 1987). Therefore, there may be a link between the evolution of non-recombining regions and sexual dimorphism.

Even when different taxa share sex chromosomes, they often differ in the size of non-recombining regions and sequence divergence between X and Y (or Z and W) chromosomes. For example, although sex chromosomes are relatively conserved in mammals, different species have different gene contents (Cortez et al., 2014). These differences may arise through differences in the timing of recombination suppression (Bergero and Charlesworth, 2009), the strength of sexually antagonistic selection (Bergero and Charlesworth, 2009; Wright et al., 2016), or population sizes (Engelstädter, 2008). However, few studies have examined whether variation in the non-recombining regions is linked to variation in sexual dimorphism. For example, previous studies on freshwater fishes belonging to the genus *Poecilia* indicate a possible association between male phenotypes and the magnitude of divergence between X and Y chromosomes (Sandkam et al., 2021; Wright et al., 2017).

A taxonomic group containing many closely related species that show diversity in both sex chromosomes and sexual dimorphism provides an excellent opportunity to investigate the link between the evolution of sex chromosomes and sexual dimorphism. The family Adrianichthyidae, commonly referred to as medaka fishes, shows frequent turnovers of sex chromosomes (Myosho et al., 2015). Since *DMY* gene was discovered as a sex-determination gene (Matsuda et al., 2002), a total of 3 sex-determining genes have been identified and 8 chromosomes have been reported to be sex-linked thus far (Myosho et al., 2015, 2012; Nagai et al., 2008; Takehana et al., 2014, 2008, 2007). Furthermore, medaka fishes, particularly in Sulawesi Island, Indonesia, show great diversity in sexual dimorphism in body coloration and fin shapes (Mokodongan and Yamahira, 2015; Sumarto et al., 2020). These fishes enable us to test whether the sex chromosome evolution is associated with the evolution of sexually dimorphic traits. Previous studies have shown that red and white pigmentation in the Japanese medaka fish *Oryzias latipes* (Aida, 1921; Wada et al., 1998) and blue coloration in the Sulawesi medaka *Oryzias woworae* (Ansai et al., 2021) are linked to their sex chromosomes. However, other sexual dimorphisms are autosomal (Ansai et al., 2021; Kawajiri et al., 2015, 2014). Therefore, it remains elusive whether sex chromosome evolution plays an important role in the evolution of sexual dimorphism in this family.

As the first step toward making the medaka fishes a model for research on the link between the evolution of sex chromosomes and sexual dimorphism, we investigated the sex chromosomes of 13 species of *Oryzias* from Sulawesi. We have three specific aims. First, we analyzed the sex chromosomes of species that either had not yet been investigated (*O. sarasinorum, O. nigrimas, O. nebulosus, O. orthognathus*, and two genetically distinct populations of *O. marmoratus*) or were recently described as a new species (*O. hadiatyae, O. eversi*, and *O. dopingdopingensis*). Second, because previous studies on the sex chromosomes of Sulawesi medaka fishes used strains that have been maintained in the laboratory for multiple generations, we analyzed sex chromosomes of wild-derived fish, except for two species. Domesticated stocks can change sex chromosomes. For example, the zebrafish has a ZW sex-determination system in the wild but lost the master sex-determining locus during domestication (Wilson et al., 2014), making the sex determination of laboratory strains polygenic (Liew et al., 2012). Finally, we investigated variation in the magnitude of divergence between the X and Y chromosomes among species that share the same sex chromosome.

## 2 MATERIAL AND METHODS

### 2.1 Fish

Thirteen species of *Oryzias* endemic to Sulawesi, Indonesia, were analyzed for their sex chromosomes (Figure 1; Table S1). For *O. marmoratus*, we used two populations: Lake Lantoa and Lake Mahalona. Specimens of two of the species, *O. eversi* and *O. sarasinorum*, were derived from a full-sib family for each species, whose parents were taken from a closed colony maintained in the lab for about 6 and 8 years, respectively. For the other species, specimens were caught from the wild. All fish were euthanized with ethyl 3-aminobenzoate methanesulfonate (MS-222, Sigma-Aldrich), and then fixed in ethanol or RNAlater solution (Thermo Fisher). Genomic DNA was isolated from muscle tissue of the fixed specimens using the DNeasy Blood & Tissue Kit (Qiagen), and the DNA concentration was determined with a Quant-iT PicoGreen dsDNA Assay Kit (Thermo Fisher).

**FIGURE 1.**
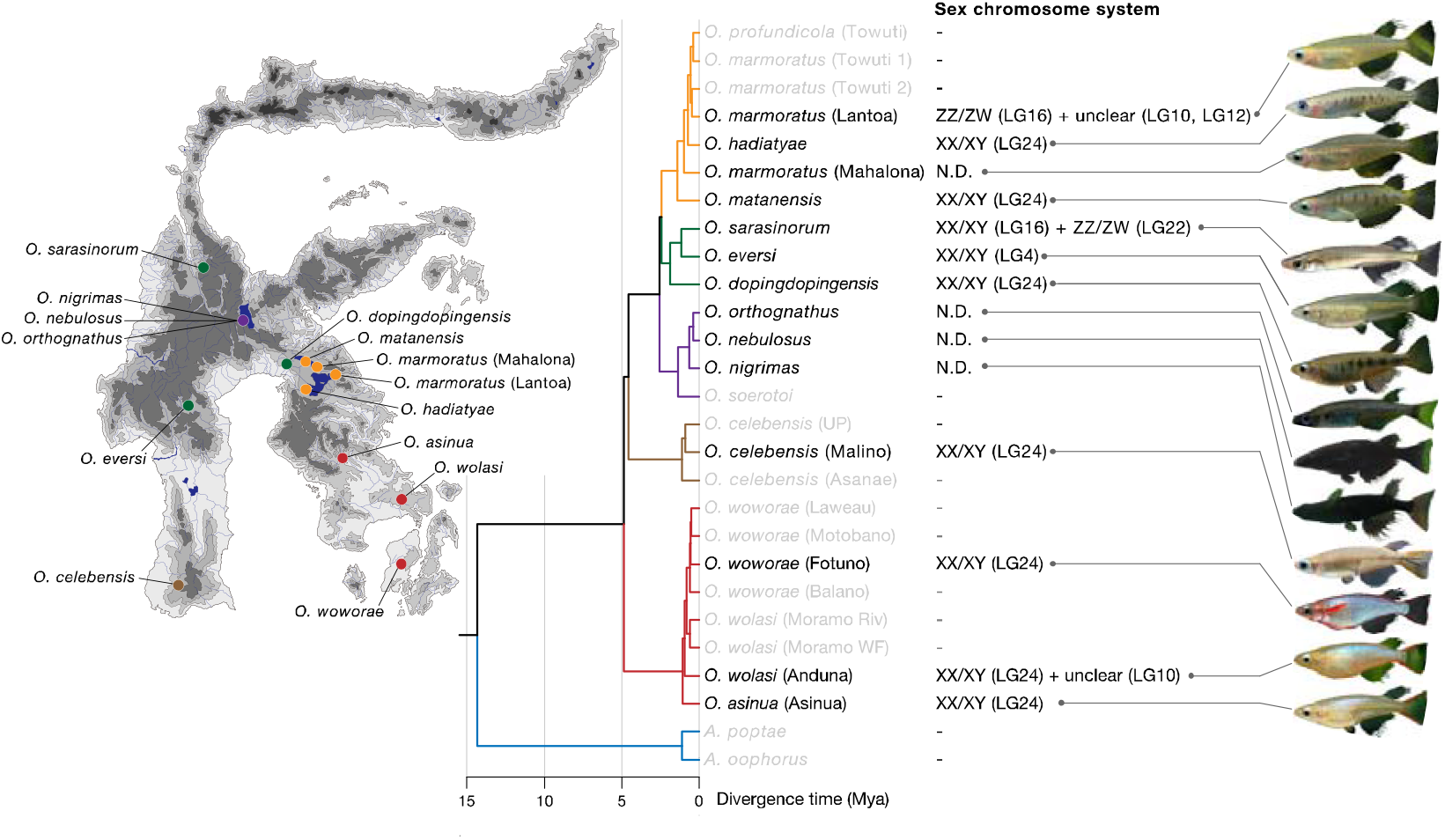
Sex-chromosome system in species of Adrianichthyidae endemic to Sulawesi, Indonesia. A map of Sulawesi showing locations of wild habitats of the 13 species analyzed in this study (map provided by Thomas von Rintelen). A chronogram shows divergence times for Sulawesi Adrianichthyidae species, which was estimated in Ansai et al. (2021). The x-axis indicates the time-scale in millions of years ago (Mya). The major lineages of Sulawesi medaka species are shown as different colors. The names of species analyzed in this study are shown in black font. Linkage groups (LG) are defined according to numbering of the chromosomes in the Japanese rice fish *Oryzias latipes*, based on genomic synteny between *O. latipes* and *O. celebensis* reference assemblies. A representative image of an adult male of each species is shown on the right side of the panel

### 2.2 Pooled sequencing

We used a pooled sequencing approach for finding the sex chromosomes (Palmer et al., 2019). For each population or species, equal molars of DNA (9–10 individuals per sex) were mixed to make a pooled DNA sample of 500 ng. Libraries were constructed using NEBNext Ultra II DNA Library Prep Kit for Illumina (NEB) with each pool indexed with a unique dual barcode (NEBNext Multiplex Oligos for Illumina, NEB). The libraries were sequenced with 150-bp paired-end reads on the Illumina HiSeq X instrument at Macrogen Japan, Tokyo, Japan. Average sequencing coverage (±SD) per pool was 34.4X (±7.8) (Table S1). The coverages of all pools exceeded a recommended sequencing depth for a heterozygosity-based approach to sex chromosome identification (Palmer et al., 2019).

The sequenced reads were trimmed using Trim Galore 0.6.4_dev with Cutadapt 2.6 and then mapped to a reference assembly of a previously constructed *Oryzias celebensis* female (OryCel_1.0) (Ansai et al., 2021) by BWA-MEM v0.7.17 with default parameters. The alignment files were sorted by genomic coordinates using *sort* in samtools v1.9, and the PCR duplicates were removed using *MarkDuplicates* in Picard Tools v2.26.9 (https://broadinstitute.github.io/picard).

Genomic differentiation between the female pool and the male pool was analyzed using PSASS v3.1.0 (https://github.com/SexGenomicsToolkit/PSASS). Briefly, a pileup file made for each sex of each population was generated using uniquely mapped reads 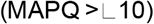. Next, the values of *F*_ST_ between two sexes and the number of sex-specific heterozygous SNPs (i.e., heterozygote in one sex while homozygous in the other sex) were calculated for each 100-kb non-overlapped window with default parameters and the group snps (--group-snps) option: in this option, consecutive SNPs are counted as one. For the subsequent analysis, windows that were not anchored to any chromosomes or have low depth of mapped reads (absolute depth < 10) were removed.

### 2.3 Identification of sex chromosomes

We first searched for windows that showed both high intersexual *F*_ST_ and high density of sex-specific heterozygous SNPs, either male-specific or female-specific. These values were judged as statistically significantly high when they exceeded the 99.9^th^ percentile of genome-wide distributions. Therefore, chromosomes with two or more candidate windows were defined as sex chromosome(s). When no sex chromosomes were identified with this criterion, we reduced the genome-wide significant threshold to the 99.5^th^ percentile (see Table S3 for the criteria used for each sex chromosome). The linkage group (LG) number of each region followed that of *O. latipes*; that is, the correspondence between the LG of *O. latipes* and the chromosome number of the reference *O. celebensis* assembly was previously determined by synteny analysis using 1-to-1 orthologous genes (Ansai et al., 2021).

To identify the sex chromosome of *O. eversi*, we additionally conducted double-digest restriction site-associated DNA sequencing (ddRAD-seq) of a lab-raised family. The ddRAD data were obtained from parents and their F_1_ offspring (25 females and 25 males), and ddRAD-seq was conducted using EcoRI and BglII, as described previously (Sutra et al., 2019). Briefly, the library was sequenced with 150-bp paired-end reads on Illumina HiSeq X at Macrogen Japan (Table S2). In total, 16,641,036 and 25,212,292 reads (2,512,796,436 and 3,807,056,092 bp) were sequenced from the G_0_ female and the G_0_ male, respectively. For the F_1_ progeny, 5,236,056 ± 1,296,817.6 reads (790,644,456 ± 195,819,458 bp) were sequenced on average (± standard deviation). The reads were trimmed and then mapped to the *Oryzias celebensis* reference assembly (OryCel_1.0) as described in Sutra et al. 2019. Variant sites were called from uniquely mapped reads 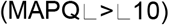 using *mpileup* and *call* in bcftools v1.5 (Li, 2011). Using VCFtools v0.1.13 (Danecek et al., 2011), we selected only SNPs with high genotyping quality 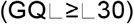, a high read depth 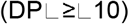, and genotyping in ≥90% of individuals in the family. We removed non-informative sites that were missing in either parent or were homozygous in both parents or showed significant segregation distortion (χ^2^ test, *P* < 0.05 with Bonferroni correction), resulting in 222 SNPs. Finally, association between sex and genotypes were tested using Fisher’s exact test with Bonferroni correction in R v4.1.0.

### 2.4 Gene ontology analysis

To investigate what kinds of genes are enriched around the sex-determining locus on LG24 in *O. hadiatyae*, we collected protein-coding genes within the region from a gene annotation dataset of the *O. celebensis* reference assembly (Ansai et al., 2021). Using BLASTN, we identified orthologs of *O. latipes* (Hd-rR strain). Using the Ensembl gene ID of *O. latipes* (Hd-rR strain), gene ontology (GO) enrichment analysis was conducted using ShinyGO v0.75 (Ge et al., 2020).

## 3 RESULTS

We first examined whether sex chromosomes previously identified in laboratory stocks were the same as those detected in our analysis using wild-caught fish. Four species (*O. celebensis, O. matanensis, O. woworae*, and *O. wolasi*) were previously reported to have XY sex-chromosome systems with LG24 being sex-linked (Myosho et al., 2015). In all of these species, LG24 showed signatures of sex chromosomes in our analysis too (Figure 2; Table S3). For example, in *O. celebensis*, three windows on LG24 showed high levels of intersexual *F*_ST_ and accumulation of male-specific heterozygous SNPs (Figure 2a; Table S3). These windows were close to marker genes that were previously reported to be sex-linked (*sox7*, chr24:14,112,606–14,115,342; *pgm3/PGM*, chr24:15,397,090–15,404,121; *hsdl1*, chr24:17,575,446–17,579,140). On LG24 of all these four species, the numbers of male-specific SNPs were elevated, indicating that these have XY systems. Interestingly, *O. wolasi* exhibited another peak on LG10 (Figure 2c; Table S3), suggesting the possible presence of a minor-effect locus in this species in the wild.

**FIGURE 2.**
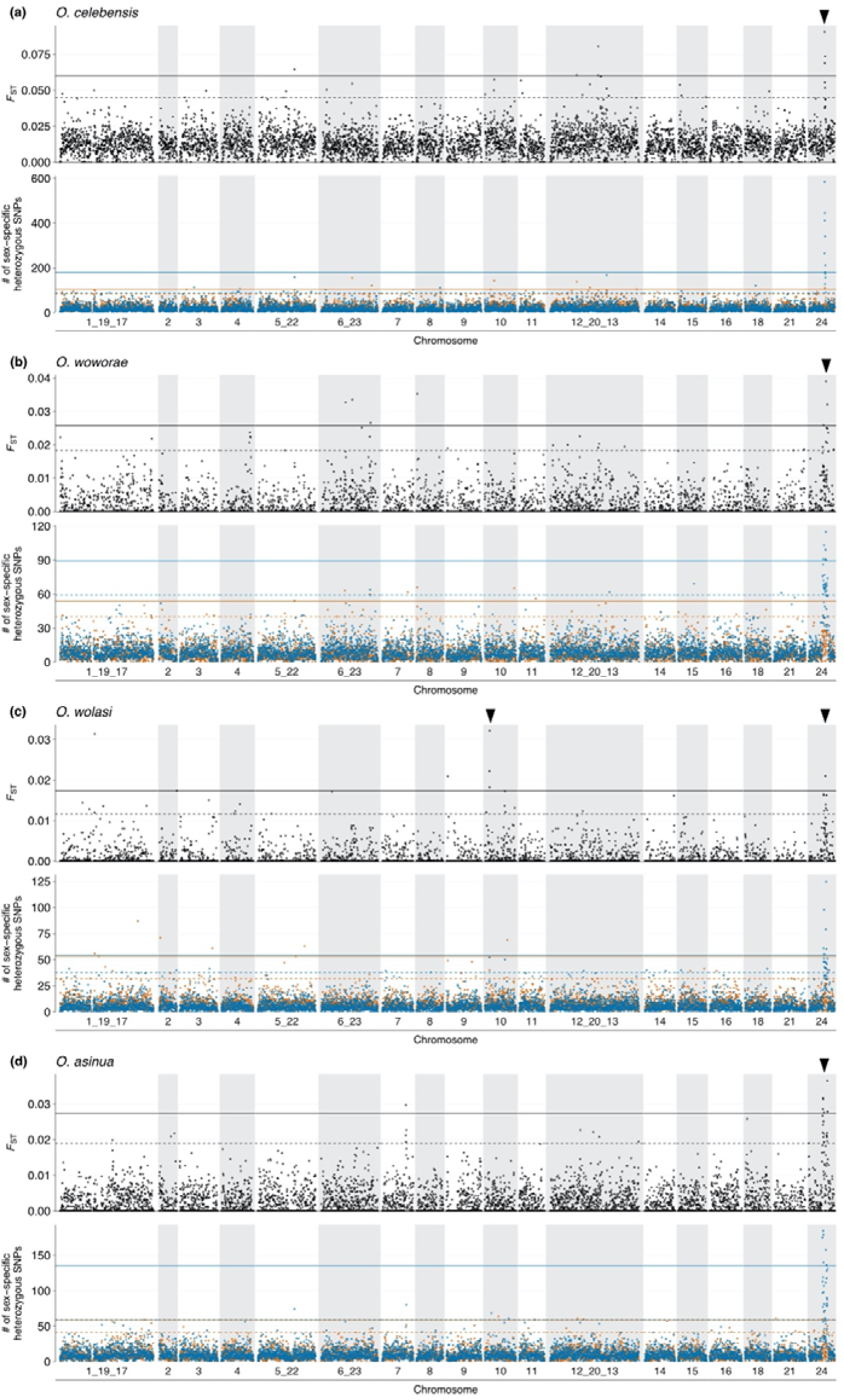
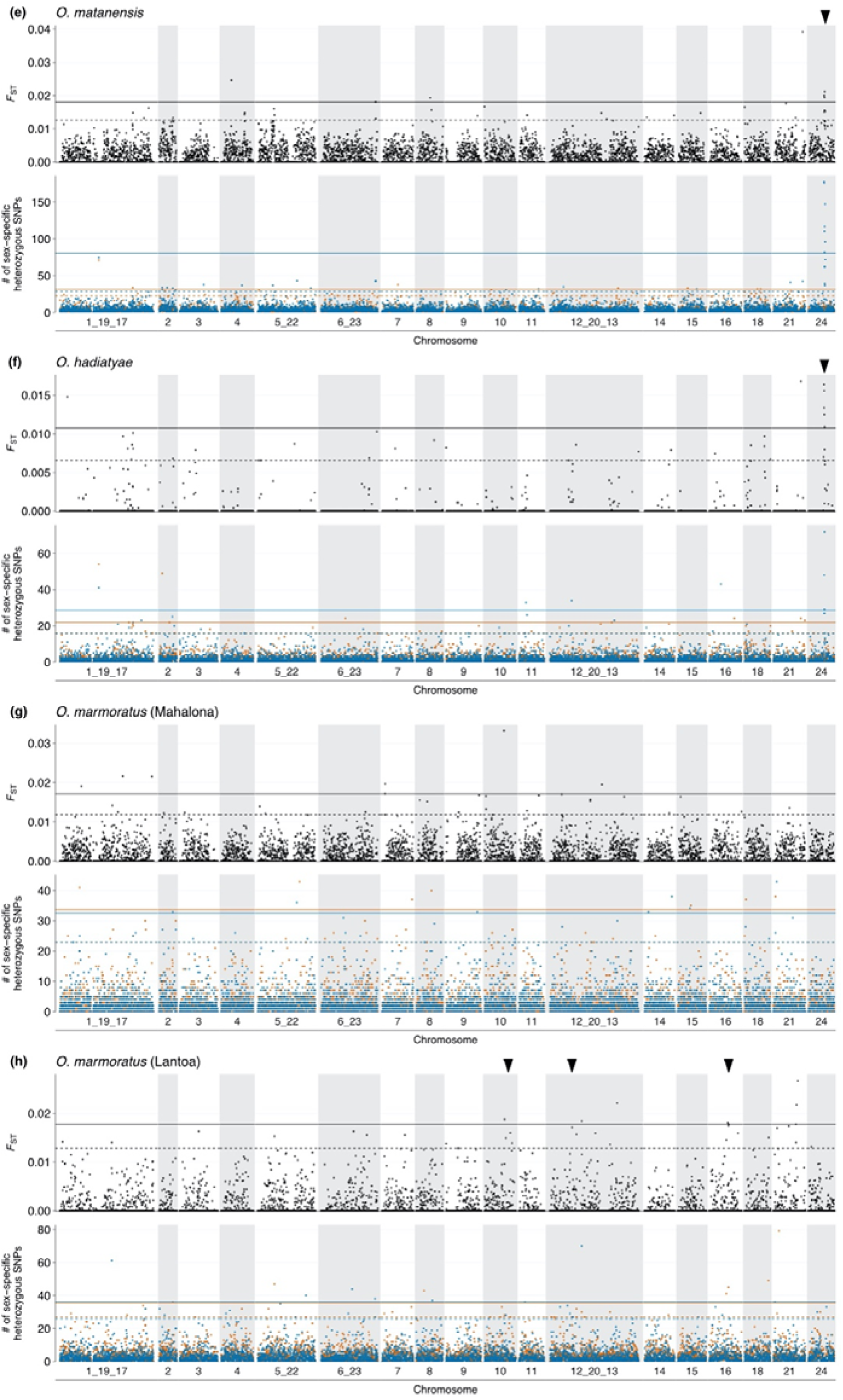
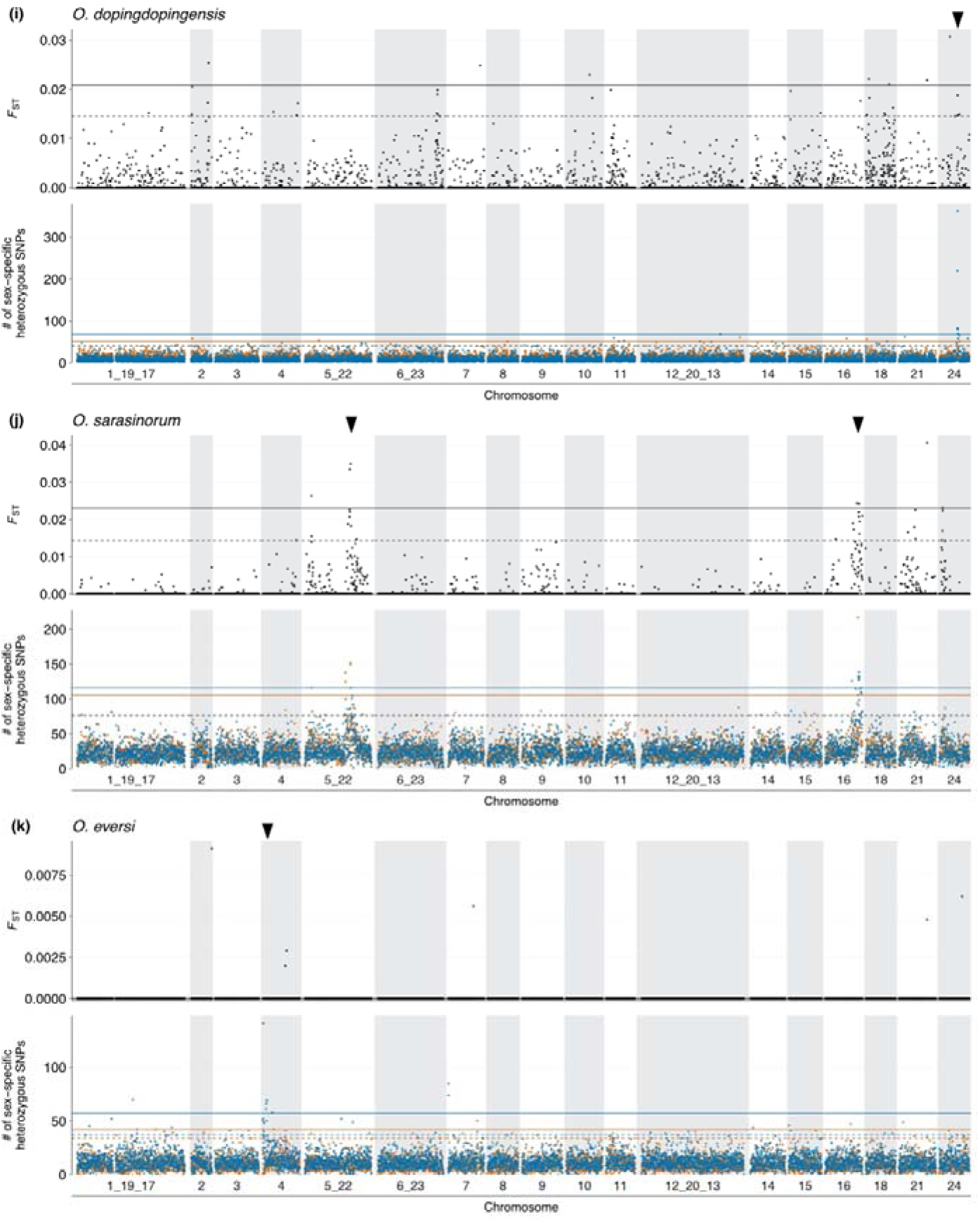
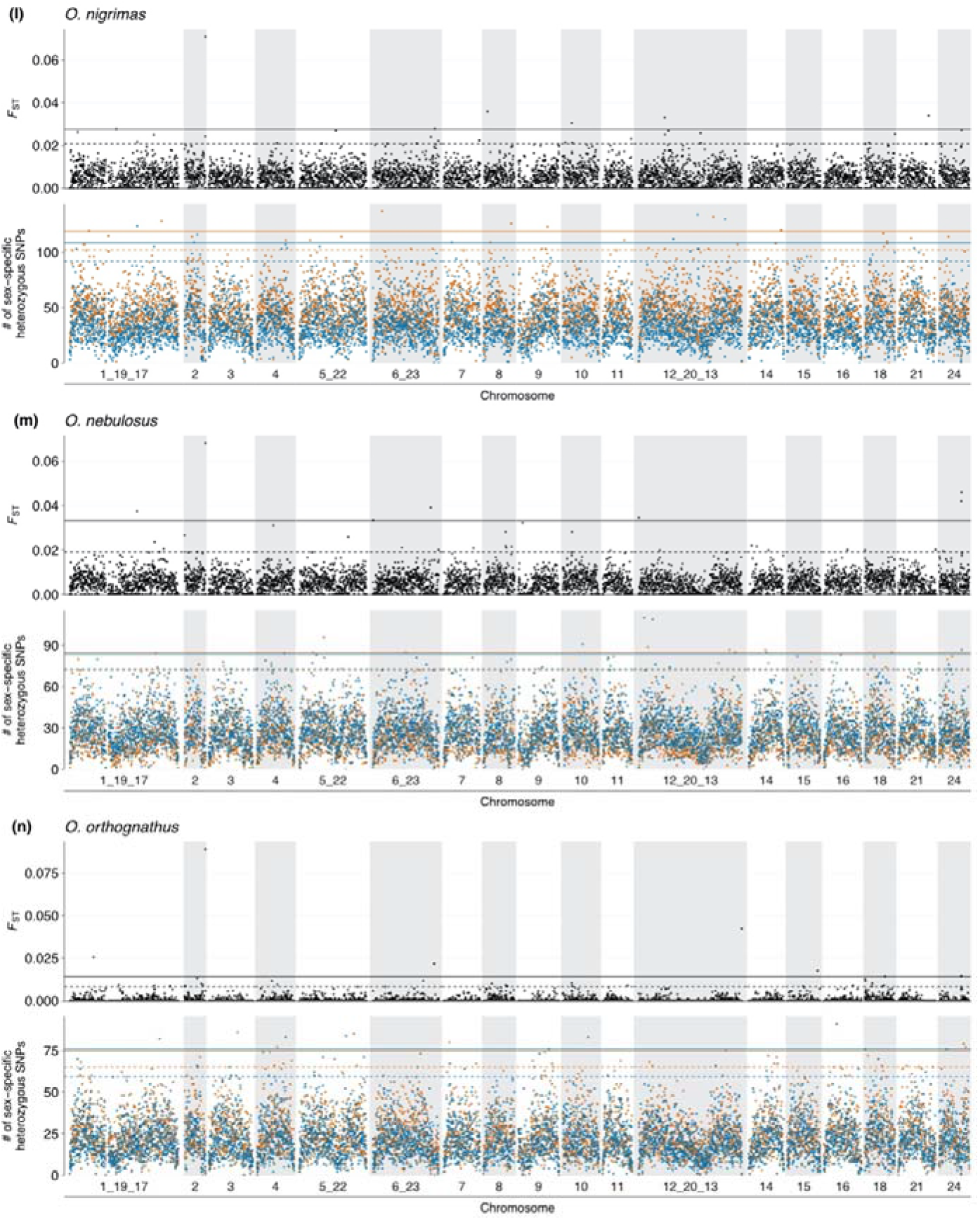
Genomic differentiation between the two sexes estimated by pooled sequencing in *Oryzias celebensis* (a), *O. woworae* (b), *O. wolasi* (c), *O. asinua* (d), *O. matanensis* (e), *O. hadiatyae* (f), *O. marmoratus* (Lake Mahalona) (g), *O. marmoratus* (Lake Lantoa) (h), *O. dopingdopingensis* (i), *O. sarasinorum* (j), *O. eversi* (k), *O. nigrimas* (l), *O. nebulosus* (m), and *O. orthognathus* (n). Intersexual *F*_ST_ and numbers of heterozygous single nucleotide polymorphisms (SNPs) in either of the sexes are calculated. Closed arrowheads indicate candidates for sex-determining loci identified in this study. Each dot shows a 100-kb non-overlapped window. Continuous and dashed lines show the 99.9^th^ percentile and 99.5^th^ percentile values, respectively. Brown and blue colors indicate numbers of heterozygous SNPs in females and males, respectively

Although a previous study showed that *O. marmoratus* in the laboratory stock derived from Lake Touwti has a XY system with LG10 sex-linked (Myosho et al., 2015), we found different results. Fish collected from Lake Lantoa showed signatures of sex chromosome on three chromosomes (LG10, LG12, and LG16). However, we did not find any signatures of sex chromosomes on any chromosomes in the *O. marmoratus* Lake Mahalona population (Figure 2g,h; Table S3). Currently, we do not know whether these differences reflect a difference between laboratory stocks and wild fish or among different geographical populations.

In addition to the previously reported four species (see above), three species newly analyzed in this study (*O. asinua, O. hadiatyae*, and *O. dopingdopingensis*) showed signatures of the XY sex chromosome on LG24 (Figure 2b-e; Table S3). The regions with signatures of sex chromosomes overlapped among these seven species, suggesting the possibility that the master sex-determining gene may be shared. These seven species with sex-linkage on LG24 enable us to investigate variation in the extent of X–Y divergence around the sex-determining loci among species. We found that they differed in the length of genomic regions with signatures of sex chromosomes. Three closely related species (*O. woworae, O. wolasi*, and *O. asinua*) had much broader regions of X–Y divergence than the other four species. Even these three species varied: *O. asinua* and *O. woworae* had more windows (27 windows in the 12.3–16.9 Mb region and the 12.4–16.9 Mb region, respectively), with excess of male-specific heterozygous SNPs than in *O. wolasi* (19 windows in the 13.2–16.7 Mb region). The other four species, *O. matanensis, O. celebensis, O. dopingdopingensis*, and *O. hadiatyae*, exhibited intersexual divergence in much more limited windows (12, 11, 10, and 6 windows, respectively). We did not find significant differences in mapped read coverage between two sexes for any of these seven species (Figure S1), suggesting that degeneration of the Y chromosome is small. *Oryzias hadiatyae* had the narrowest region with signatures of sex chromosomes within a 800-kb region (14.1–14.9 Mb) on LG 24. This region includes 31 genes according to the annotation of *O. latipes* Ensembl. No significant enrichment for any GO terms was found.

New sex chromosomes were identified in two species. *Oryzias sarasinorum* showed sex chromosome signatures on LG16 and LG22 (Figure 2j). Whereas 4 outlier windows at 25.9–26.4 Mb on LG16 were enriched for male-specific heterozygous SNPs, 3 windows at 34.6–35.4 Mb of LG22 were enriched for female-specific heterozygous SNPs (Figure 4a, b; Table S3). These results suggest the possibility that this species may have a mixture of XY and ZW systems, as observed in cichlids (Roberts et al., 2016). *Oryzias eversi*, which diverged from *O. sarasinurum* at approximately 170,000 years ago (Horoiwa et al., 2021), did not show any significant signatures of sex chromosomes based on our criteria (Figure 2k); however, male-specific heterozygous SNPs tended to accumulate in 5 windows located at the 5’-end of LG4 (i.e., 0.3, 2.5, 2.7, 3.1, and 7.4 Mb; >99.9^th^ percentile) (Figure 4c). To test the sex-linkage of this locus, we genotyped a lab-raised family with ddRAD-seq. We found three SNPs significantly associated with sex (chr4:141,042, chr4:8,924,730, and chr9,530,418; *P* < 0.05, Fisher’s exact test with Bonferroni correction). One of them on LG4 (chr4:141,042) was perfectly associated with sex, with all 25 males being heterozygote and all 25 females being homozygote (Table S4). These results suggest that LG4 may be XY chromosomes in *O. eversi*.

**FIGURE 3.**
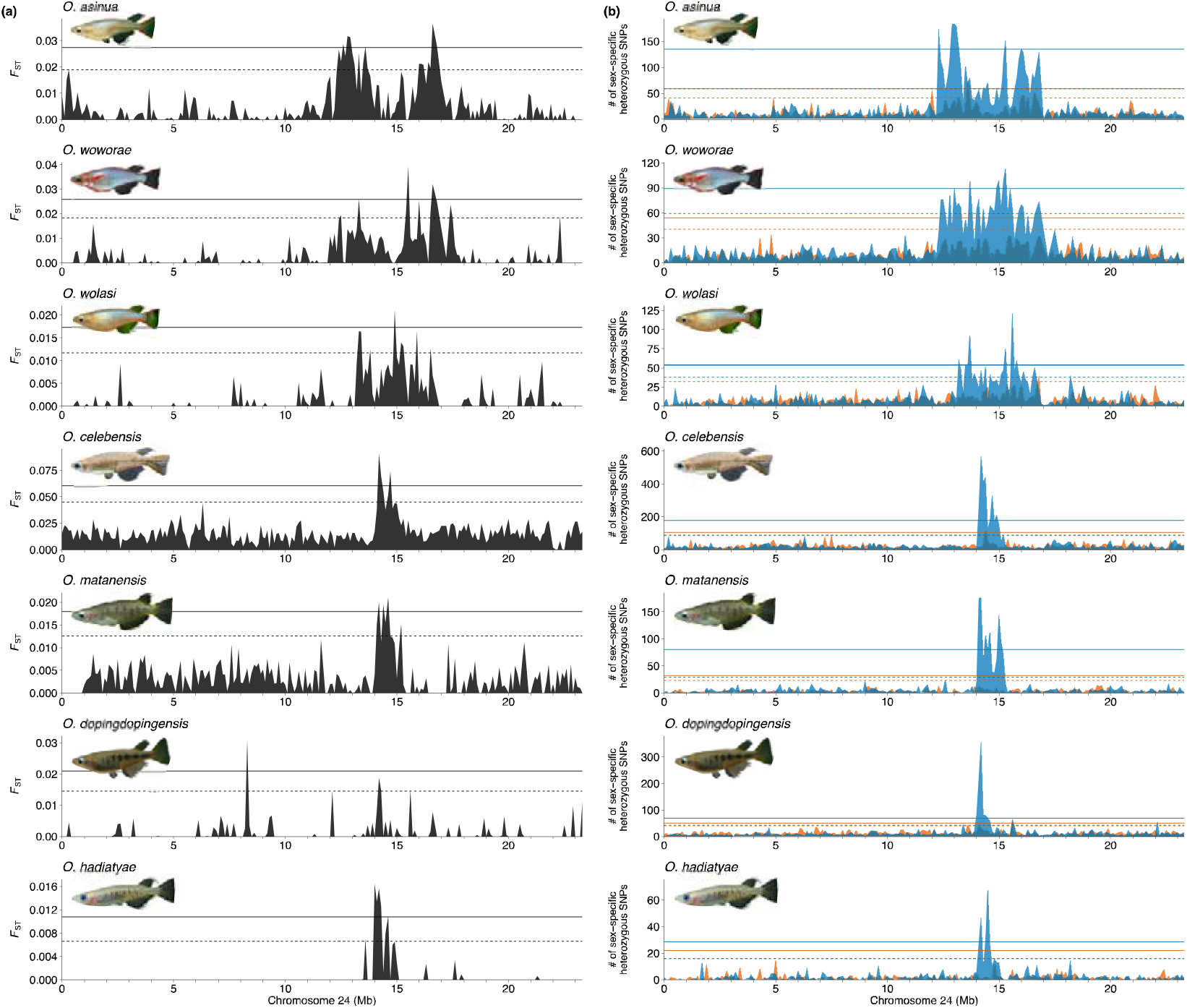
Intersexual differentiation on chromosome 24, which is a sex chromosome in 7 *Oryzias* species. Intersexual *F*_ST_ (a), and numbers of heterozygous single nucleotide polymorphisms (SNPs) in either of two sexes (b), are shown in each 100 kb of non-overlapped window. Continuous and dashed lines denote the 99.9^th^ percentile and 99.5^th^ percentiles, respectively. Brown and blue colors indicate numbers of heterozygous SNPs in females and males, respectively

**FIGURE 4.**
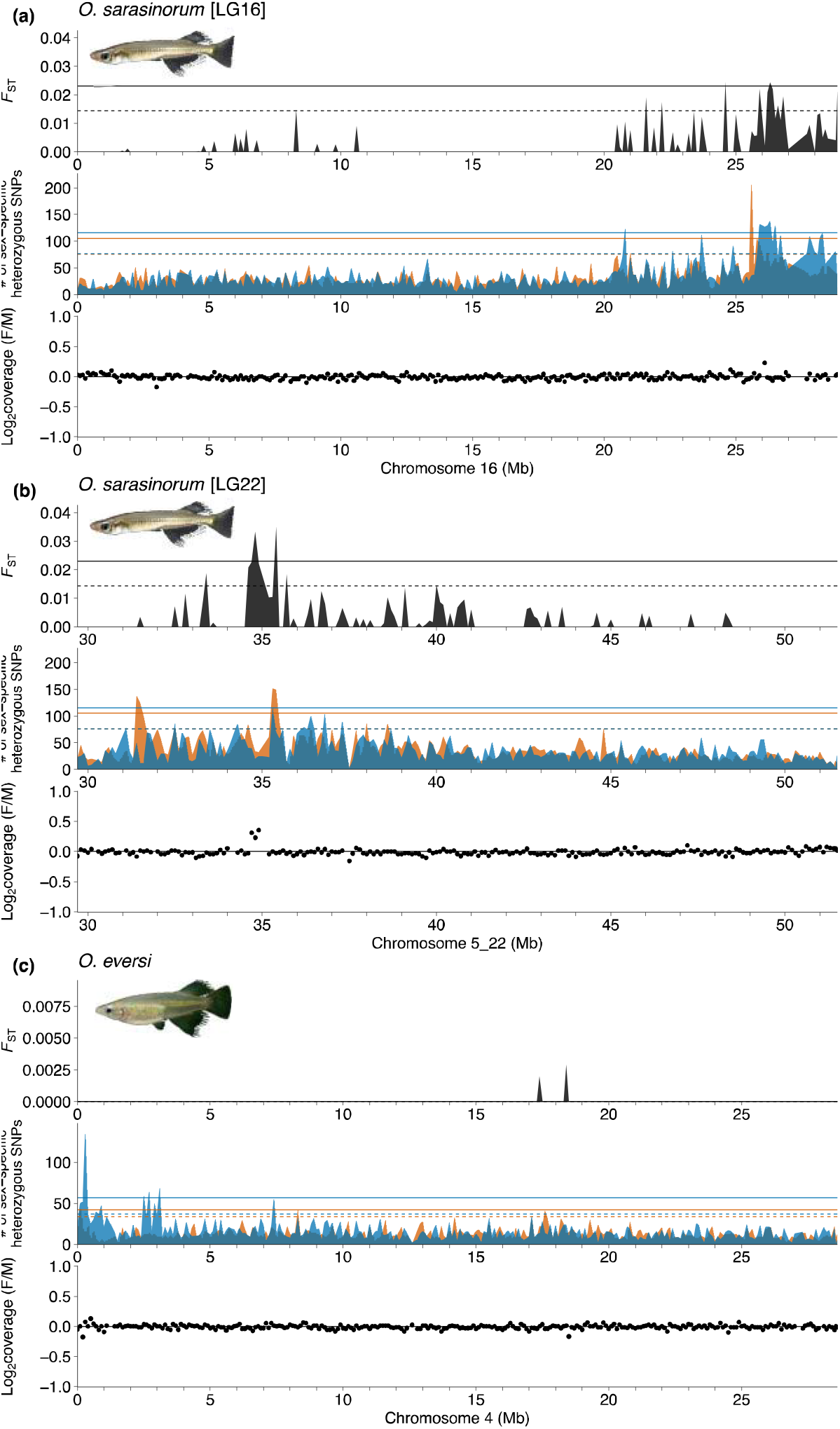
Intersexual differentiation on a sex chromosome of *Oryzias eversi* (a) and *O. sarasinorum* (b, c). Intersexual *F*_ST_, numbers of heterozygous single nucleotide polymorphisms (SNPs) in either of the sexes (b), and read coverage Log_2_ (female reads/male reads) are shown in each 100 kb of non-overlapped window. Continuous and dashed lines show the 99.9^th^ percentile and 99.5^th^ percentile values, respectively. Brown and blue colors indicate numbers of heterozygous SNPs in females and males, respectively

The three species endemic to Lake Poso (*O. nigrimas, O. nebulosus*, and *O. orthognathus*) did not exhibit any significant signatures from sex chromosomes on any chromosome (Figure 2l–n).

## 4 DISCUSSION

We found great diversity in sex chromosomes among Sulawesi medaka species, both in the linkage groups associated with sex and the magnitude of differentiation between X and Y sex chromosomes. LG24 is the most frequently observed linkage group associated with sex, with seven of the medaka species using LG24 as XY sex chromosomes. These seven species belong to 4 of 6 different major clades of Sulawesi medaka fishes, and their geographic ranges on the island are quite distant (apart) from each other (Figure 1). There are two possibilities for the origin of LG24 sex chromosomes: these sex chromosome evolved once at an early stage of Sulawesi medaka diversification or they evolved multiple times. Although we cannot exclude either possibility, the patterns of divergence between X and Y seem to be similar among the monophyletic group *O. woworae, O. wolasi*, and *O. asinua* (Figure 3), suggesting that the sex chromosomes of this clade may have the same origin. Identification of sex-determining mutations is a possibility in other medaka species (Matsuda et al., 2002; Myosho et al., 2015, 2012; Takehana et al., 2014), and verification would help to answer which scenario, the single or multiple origins of LG24 sex chromosomes, is more likely.

The seven species with sex-linked LG24 differed in the length and magnitude of X–Y divergence. One possible mechanism for different divergence levels is the different timing of the evolution of recombination suppression. In several species, such as among mammals and sticklebacks, there are strata on sex chromosomes, which is thought to have arisen through multiple events of chromosomal rearrangements on the sex chromosomes. Further comparative analysis of chromosomal structures and recombination rates is necessary to test this possibility. Sexually antagonistic selection can promote the evolution of recombination suppression and X–Y divergence (Bergero and Charlesworth, 2009; Rice, 1987; Wright et al., 2016). Our previous analysis revealed that a quantitative trait locus for blue coloration on the lateral body surface, which is a sexually dimorphic trait found in *O. woworae* males, is located on LG24 (Ansai et al., 2021). Given that the blue body coloration is shared by *O. wolasi* and *O. asinua* and that these three species have larger regions of X–Y divergence than the other four species known to possess the sex-linked LG24, it is possible that sexually antagonistic selection could play a role in X–Y divergence. However, it should be noted that red coloration, another sexually dimorphic trait, was mapped on autosome (Ansai et al., 2021). Further identification of loci responsible for potential sexually antagonistic traits in Sulawesi medaka species would help us test the link between the evolution of sexual dimorphism and sex chromosomes.

This study identified new sex chromosomes in two closely related species: *O. sarasinorum* and *O. eversi*. In *O. sarasinorum*, LG16 and LG22 are associated with sex. Although a previous study showed that LG16 is sex-linked in *O. javanicus* (Takehana et al., 2008), the sex-determining regions did not overlap. Interestingly, the divergent window (25.9–26.4 Mb) on LG16 contains genes located on the SD locus of Atlantic cod *Gadus morhua (cep76*, chr16:25,931,989–25,950,574; *ptpn2a*, chr16:25,953,059–25,965,311) (Star et al., 2016). A recent study demonstrated that the Y-specific sequence of Atlantic cod contains a single gene named *zkY* (Kirubakaran et al., 2019), suggesting that investigating whether a similar gene is located on the Y sequence of *O. sarasinorum* is promising. The sex chromosome of *O. eversi* was found to be LG4, which has not been reported as a sex chromosome in any other medaka species. In *O. eversi*, a SNP at the 5’ end of LG4 (chr4:141,042) was perfectly associated with sex. Near this locus (chr4:219,486–226,125), there exists the *amh* gene, which encodes anti-Müllerian hormone that regulates gonad development and germ-cell proliferation in medaka fishes (Klüver et al., 2007; Morinaga et al., 2007; Nakamura et al., 2012). Recent studies have demonstrated that *amh* has repeatedly evolved as an SD gene in different teleost fishes, including the Patagonian pejerrey *Odontesthes hatcheri* (Hattori et al., 2012), Northern pike *Esox lucius* (Pan et al., 2019), threespine stickleback *Gasterosteus aculeatus* (Peichel et al., 2020), and *Sebastes* rockfish (Song et al., 2021). Further studies are needed to identify causal mutations on the Y allele and to analyze genetic diversity of the sex chromosomes in wild populations.

We found that sex chromosome systems in two wild populations of *O. marmoratus*, collected from Lake Mahalona and Lake Lantoa, differ from that reported for a laboratory stock originally derived from Lake Towuti (Myosho et al., 2015). The first possible explanation for the discrepancy is geographical variation. Our recent study using genome-wide SNPs demonstrated that wild populations of *O. marmoratus* are not monophyletic (Mandagi et al., 2021). Therefore, a taxonomic species classified as *O. marmoratus* may be composed of several genetically distinct species. Another possibility is that domestication changed the sex chromosomes, as observed in the zebrafish (Liew et al., 2012; Wilson et al., 2014). Analysis of sex chromosomes in natural populations of Lake Towuti *O. marmoratus* is necessary to tell which theory is correct.

No apparent signatures of sex chromosomes were found in *Oryzias* species endemic to Lake Poso (*O. nigrimas, O. nebulosus*, and *O. orthognathus*). Currently, we are unsure whether these species lack sex chromosomes. Alternatively, it is possible that they have multiple sex chromosomes, but with each one having a small effect, and the small sample sizes in this study precluded the identification of such sex chromosomes. Genome-wide association studies using whole-genome sequencing datasets of more individuals may enable the identification of weak and/or polygenic signatures of the sex chromosomes.

Although theoretical studies predict that sex chromosome evolution can drive the evolution of sexual dimorphism, empirical studies are still limited (Dean and Mank, 2014; Kitano and Peichel, 2012; Rice, 1984). A taxonomic group that shows diversity in both sex chromosomes and sexual dimorphism can provide us with excellent opportunities to investigate the link between the evolution of sex chromosomes and sexual dimorphism. Our study demonstrates that frequent turnovers and the great diversity of the sex chromosomes make the Sulawesian medaka a model system for investigating the roles of sex chromosome evolution in the diversification of sexual dimorphism.

## ACKNOWLEDGEMENTS

We thank the Ministry of Research, Technology, and Higher Education, Republic of Indonesia (RISTEKDIKTI) and the Faculty of Fisheries and Marine Science, Sam Ratulangi University, for permission to conduct research in Sulawesi (Research Permit Numbers 394/SIP/FRP/SM/XI/2014, 397/SIP/FRP/SM/XI/2014, and 106/SIP/FRP/E5/Dit.KI/IV/2018). We also thank S. Kondo (Ryukoku University) and members of the Yamahira lab at the University of the Ryukyus for technical assistance. Cynthia Kulongowski with Edanz (https://jp.edanz.com/ac) edited the language of a draft of this manuscript. This work was supported by JSPS KAKENHI Grant Numbers 21K15143 to S.A., 19K16203 to J.M., and 21H04782 to K.Y.; and JST CREST Grant Number JPMJCR20S2 to J.K., K.Y., and S.A.

## AUTHOR CONTRIBUTIONS

S.A., K.Y., and J.K. conceived and designed the research. S.A., J.M., A.J.N., K.Y., and J.K. performed the experiments. S.A. analyzed the data. K.W.A.M. contributed materials. S.A. and J.K. wrote the manuscript. All authors have read and approved the final manuscript.

## DATA AVAILABILITY STATEMENT

All sequence reads will be available from the DDBJ Sequence Read Archive (SRA): short sequence reads for pool sequencing (DRAXXXXXX) and ddRAD-seq reads for a family of *Oryzias eversi* (DRAXXXXXX).

## REFERENCES

Aida, T., 1921. On the inheritance of color in a fresh-water fish, *Aplocheilus latipes* Temmick and Schlegel, with special reference to sex-linked inheritance. Genetics 6, 554–573. https://doi.org/10.1093/genetics/6.6.554

Ansai, S., Mochida, K., Fujimoto, S., Mokodongan, D.F., Sumarto, B.K.A., Masengi, K.W.A., Hadiaty, R.K., Nagano, A.J., Toyoda, A., Naruse, K., Yamahira, K., Kitano, J., 2021. Genome editing reveals fitness effects of a gene for sexual dichromatism in Sulawesian fishes. Nat. Commun. 12, 1350. https://doi.org/10.1038/s41467-021-21697-0

Bachtrog, D., Mank, J.E., Peichel, C.L., Kirkpatrick, M., Otto, S.P., Ashman, T.-L., Hahn, M.W., Kitano, J., Mayrose, I., Ming, R., Perrin, N., Ross, L., Valenzuela, N., Vamosi, J.C., 2014. Sex Determination: Why So Many Ways of Doing It? PLOS Biol. 12, e1001899. https://doi.org/10.1371/journal.pbio.1001899

Bergero, R., Charlesworth, D., 2009. The evolution of restricted recombination in sex chromosomes. Trends Ecol. Evol. 24, 94–102. https://doi.org/10.1016/j.tree.2008.09.010

Beukeboom, L.W., Perrin, N., 2014. The Evolution of Sex Determination. Oxford University Press, Oxford. https://doi.org/10.1093/acprof:oso/9780199657148.001.0001

Charlesworth, D., Charlesworth, B., 1980. Sex differences in fitness and selection for centric fusions between sex-chromosomes and autosomes. Genet. Res. 35, 205–214. https://doi.org/10.1017/S0016672300014051

Cortez, D., Marin, R., Toledo-Flores, D., Froidevaux, L., Liechti, A., Waters, P.D., Grützner, F., Kaessmann, H., 2014. Origins and functional evolution of Y chromosomes across mammals. Nature 508, 488–493. https://doi.org/10.1038/nature13151

Danecek, P., Auton, A., Abecasis, G., Albers, C.A., Banks, E., DePristo, M.A., Handsaker, R.E., Lunter, G., Marth, G.T., Sherry, S.T., McVean, G., Durbin, R., 2011. The variant call format and VCFtools. Bioinformatics 27, 2156–2158. https://doi.org/10.1093/bioinformatics/btr330

Dean, R., Mank, J.E., 2014. The role of sex chromosomes in sexual dimorphism: discordance between molecular and phenotypic data. J. Evol. Biol. 27, 1443–1453. https://doi.org/10.1111/jeb.12345

Engelstädter, J., 2008. Muller’s Ratchet and the Degeneration of Y Chromosomes: A Simulation Study. Genetics 180, 957–967. https://doi.org/10.1534/genetics.108.092379

Gammerdinger, W.J., Kocher, T.D., 2018. Unusual Diversity of Sex Chromosomes in African Cichlid Fishes. Genes 9, 480. https://doi.org/10.3390/genes9100480

Ge, S.X., Jung, D., Yao, R., 2020. ShinyGO: a graphical gene-set enrichment tool for animals and plants. Bioinformatics 36, 2628–2629. https://doi.org/10.1093/bioinformatics/btz931

Hattori, R.S., Murai, Y., Oura, M., Masuda, S., Majhi, S.K., Sakamoto, T., Fernandino, J.I., Somoza, G.M., Yokota, M., Strüssmann, C.A., 2012. A Y-linked anti-Müllerian hormone duplication takes over a critical role in sex determination. Proc. Natl. Acad. Sci. 109, 2955–2959. https://doi.org/10.1073/pnas.1018392109

Horoiwa, M., Mandagi, I.F., Sutra, N., Montenegro, J., Tantu, F.Y., Masengi, K.W.A., Nagano, A.J., Kusumi, J., Yasuda, N., Yamahira, K., 2021. Mitochondrial introgression by ancient admixture between two distant lacustrine fishes in Sulawesi Island. PLOS ONE 16, e0245316. https://doi.org/10.1371/journal.pone.0245316

Jeffries, D.L., Gerchen, J.F., Scharmann, M., Pannell, J.R., 2021. A neutral model for the loss of recombination on sex chromosomes. Philos. Trans. R. Soc. B Biol. Sci. 376, 20200096. https://doi.org/10.1098/rstb.2020.0096

Kawajiri, M., Fujimoto, S., Yoshida, K., Yamahira, K., Kitano, J., 2015. Genetic Architecture of the Variation in Male-Specific Ossified Processes on the Anal Fins of Japanese Medaka. G3 GenesGenomesGenetics 5, 2875–2884. https://doi.org/10.1534/g3.115.021956

Kawajiri, M., Yoshida, K., Fujimoto, S., Mokodongan, D.F., Ravinet, M., Kirkpatrick, M., Yamahira, K., Kitano, J., 2014. Ontogenetic stage-specific quantitative trait loci contribute to divergence in developmental trajectories of sexually dimorphic fins between medaka populations. Mol. Ecol. 23, 5258–5275. https://doi.org/10.1111/mec.12933

Kirubakaran, T.G., Andersen, Ø., De Rosa, M.C., Andersstuen, T., Hallan, K., Kent, M.P., Lien, S., 2019. Characterization of a male specific region containing a candidate sex determining gene in Atlantic cod. Sci. Rep. 9, 116. https://doi.org/10.1038/s41598-018-36748-8

Kitano, J., Peichel, C.L., 2012. Turnover of sex chromosomes and speciation in fishes. Environ. Biol. Fishes 94, 549–558. https://doi.org/10.1007/s10641-011-9853-8

Kitano, J., Ross, J.A., Mori, S., Kume, M., Jones, F.C., Chan, Y.F., Absher, D.M., Grimwood, J., Schmutz, J., Myers, R.M., Kingsley, D.M., Peichel, C.L., 2009. A role for a neo-sex chromosome in stickleback speciation. Nature 461, 1079–1083. https://doi.org/10.1038/nature08441

Klüver, N., Pfennig, F., Pala, I., Storch, K., Schlieder, M., Froschauer, A., Gutzeit, H.O., Schartl, M., 2007. Differential expression of anti-Müllerian hormone (amh) and anti-Müllerian hormone receptor type II (amhrII) in the teleost medaka. Dev. Dyn. 236, 271–281. https://doi.org/10.1002/dvdy.20997

Lenormand, T., Roze, D., 2022. Y recombination arrest and degeneration in the absence of sexual dimorphism. Science 375, 663–666. https://doi.org/10.1126/science.abj1813

Li, H., 2011. A statistical framework for SNP calling, mutation discovery, association mapping and population genetical parameter estimation from sequencing data. Bioinformatics 27, 2987–2993. https://doi.org/10.1093/bioinformatics/btr509

Liew, W.C., Bartfai, R., Lim, Z., Sreenivasan, R., Siegfried, K.R., Orban, L., 2012. Polygenic Sex Determination System in Zebrafish. PLOS ONE 7, e34397. https://doi.org/10.1371/journal.pone.0034397

Mandagi, I.F., Kakioka, R., Montenegro, J., Kobayashi, H., Masengi, K.W.A., Inomata, N., Nagano, A.J., Toyoda, A., Ansai, S., Matsunami, M., Kimura, R., Kitano, J., Kusumi, J., Yamahira, K., 2021. Species divergence and repeated ancient hybridization in a Sulawesian lake system. J. Evol. Biol. 34, 1767–1780. https://doi.org/10.1111/jeb.13932

Mank, J.E., 2009. Sex Chromosomes and the Evolution of Sexual Dimorphism: Lessons from the Genome. Am. Nat. 173, 141–150. https://doi.org/10.1086/595754

Matsuda, M., Nagahama, Y., Shinomiya, A., Sato, T., Matsuda, C., Kobayashi, T., Morrey, C.E., Shibata, N., Asakawa, S., Shimizu, N., Hori, H., Hamaguchi, S., Sakaizumi, M., 2002. DMY is a Y-specific DM-domain gene required for male development in the medaka fish. Nature 417, 559–563. https://doi.org/10.1038/nature751

Miura, I., 2017. Sex Determination and Sex Chromosomes in Amphibia. Sex. Dev. 11, 298–306. https://doi.org/10.1159/000485270

Mokodongan, D.F., Yamahira, K., 2015. Origin and intra-island diversification of Sulawesi endemic Adrianichthyidae. Mol. Phylogenet. Evol. 93, 150–160. https://doi.org/10.1016/j.ympev.2015.07.024

Morinaga, C., Saito, D., Nakamura, S., Sasaki, T., Asakawa, S., Shimizu, N., Mitani, H., Furutani-Seiki, M., Tanaka, M., Kondoh, H., 2007. The hotei mutation of medaka in the anti-Müllerian hormone receptor causes the dysregulation of germ cell and sexual development. Proc. Natl. Acad. Sci. 104, 9691–9696. https://doi.org/10.1073/pnas.0611379104

Myosho, T., Otake, H., Masuyama, H., Matsuda, M., Kuroki, Y., Fujiyama, A., Naruse, K., Hamaguchi, S., Sakaizumi, M., 2012. Tracing the Emergence of a Novel Sex-Determining Gene in Medaka, Oryzias luzonensis. Genetics 191, 163–170. https://doi.org/10.1534/genetics.111.137497

Myosho, T., Takehana, Y., Hamaguchi, S., Sakaizumi, M., 2015. Turnover of Sex Chromosomes in Celebensis Group Medaka Fishes. G3 GenesGenomesGenetics 5, 2685–2691. https://doi.org/10.1534/g3.115.021543

Nagai, T., Takehana, Y., Hamaguchi, S., Sakaizumi, M., 2008. Identification of the sex-determining locus in the Thai medaka, Oryzias minutillus. Cytogenet. Genome Res. 121, 137–142. https://doi.org/10.1159/000125839

Nakamura, S., Watakabe, I., Nishimura, T., Picard, J.-Y., Toyoda, A., Taniguchi, Y., di Clemente, N., Tanaka, M., 2012. Hyperproliferation of mitotically active germ cells due to defective anti-Müllerian hormone signaling mediates sex reversal in medaka. Development 139, 2283–2287. https://doi.org/10.1242/dev.076307

Palmer, D.H., Rogers, T.F., Dean, R., Wright, A.E., 2019. How to identify sex chromosomes and their turnover. Mol. Ecol. 28, 4709–4724. https://doi.org/10.1111/mec.15245

Pan, Q., Feron, R., Yano, A., Guyomard, R., Jouanno, E., Vigouroux, E., Wen, M., Busnel, J.-M., Bobe, J., Concordet, J.-P., Parrinello, H., Journot, L., Klopp, C., Lluch, J., Roques, C., Postlethwait, J., Schartl, M., Herpin, A., Guiguen, Y., 2019. Identification of the master sex determining gene in Northern pike (Esox lucius) reveals restricted sex chromosome differentiation. PLOS Genet. 15, e1008013. https://doi.org/10.1371/journal.pgen.1008013

Peichel, C.L., McCann, S.R., Ross, J.A., Naftaly, A.F.S., Urton, J.R., Cech, J.N., Grimwood, J., Schmutz, J., Myers, R.M., Kingsley, D.M., White, M.A., 2020. Assembly of the threespine stickleback Y chromosome reveals convergent signatures of sex chromosome evolution. Genome Biol. 21, 177. https://doi.org/10.1186/s13059-020-02097-x

Ponnikas, S., Sigeman, H., Abbott, J.K., Hansson, B., 2018. Why Do Sex Chromosomes Stop Recombining? Trends Genet. 34, 492–503. https://doi.org/10.1016/j.tig.2018.04.001

Rice, W.R., 1987. The Accumulation of Sexually Antagonistic Genes as a Selective Agent Promoting the Evolution of Reduced Recombination between Primitive Sex Chromosomes. Evolution 41, 911–914. https://doi.org/10.2307/2408899

Rice, W.R., 1984. Sex Chromosomes and the Evolution of Sexual Dimorphism. Evolution 38, 735–742. https://doi.org/10.2307/2408385

Roberts, N.B., Juntti, S.A., Coyle, K.P., Dumont, B.L., Stanley, M.K., Ryan, A.Q., Fernald, R.D., Roberts, R.B., 2016. Polygenic sex determination in the cichlid fish Astatotilapia burtoni. BMC Genomics 17, 835. https://doi.org/10.1186/s12864-016-3177-1

Roberts, R.B., Ser, J.R., Kocher, T.D., 2009. Sexual Conflict Resolved by Invasion of a Novel Sex Determiner in Lake Malawi Cichlid Fishes. Science 326, 998–1001. https://doi.org/10.1126/science.1174705

Ross, J.A., Urton, J.R., Boland, J., Shapiro, M.D., Peichel, C.L., 2009. Turnover of Sex Chromosomes in the Stickleback Fishes (Gasterosteidae). PLOS Genet. 5, e1000391. https://doi.org/10.1371/journal.pgen.1000391

Sandkam, B.A., Almeida, P., Darolti, I., Furman, B.L.S., van der Bijl, W., Morris, J., Bourne, G.R., Breden, F., Mank, J.E., 2021. Extreme Y chromosome polymorphism corresponds to five male reproductive morphs of a freshwater fish. Nat. Ecol. Evol. 5, 939–948. https://doi.org/10.1038/s41559-021-01452-w

Song, W., Xie, Y., Sun, M., Li, X., Fitzpatrick, C.K., Vaux, F., O’Malley, K.G., Zhang, Q., Qi, J., He, Y., 2021. A duplicated *amh* is the master sex-determining gene for *Sebastes* rockfish in the Northwest Pacific. Open Biol. 11, 210063. https://doi.org/10.1098/rsob.210063

Star, B., Tørresen, O.K., Nederbragt, A.J., Jakobsen, K.S., Pampoulie, C., Jentoft, S., 2016. Genomic characterization of the Atlantic cod sex-locus. Sci. Rep. 6, 31235. https://doi.org/10.1038/srep31235

Sumarto, B.K.A., Kobayashi, H., Kakioka, R., Tanaka, R., Maeda, K., Tran, H.D., Koizumi, N., Morioka, S., Bounsong, V., Watanabe, K., Musikasinthorn, P., Tun, S., Yun, L.K.C., Anoop, V.K., Raghavan, R., Masengi, K.W.A., Fujimoto, S., Yamahira, K., 2020. Latitudinal variation in sexual dimorphism in a freshwater fish group. Biol. J. Linn. Soc. 131, 898–908. https://doi.org/10.1093/biolinnean/blaa166

Sutra, N., Kusumi, J., Montenegro, J., Kobayashi, H., Fujimoto, S., Masengi, K.W.A., Nagano, A.J., Toyoda, A., Matsunami, M., Kimura, R., Yamahira, K., 2019. Evidence for sympatric speciation in a Wallacean ancient lake. Evolution 73, 1898–1915. https://doi.org/10.1111/evo.13821

Takehana, Y., Hamaguchi, S., Sakaizumi, M., 2008. Different origins of ZZ/ZW sex chromosomes in closely related medaka fishes, Oryzias javanicus and O. hubbsi. Chromosome Res. 16, 801–811. https://doi.org/10.1007/s10577-008-1227-5

Takehana, Y., Matsuda, M., Myosho, T., Suster, M.L., Kawakami, K., Shin-I, T., Kohara, Y., Kuroki, Y., Toyoda, A., Fujiyama, A., Hamaguchi, S., Sakaizumi, M., Naruse, K., 2014. Co-option of Sox3 as the male-determining factor on the Y chromosome in the fish Oryzias dancena. Nat. Commun. 5, 4157. https://doi.org/10.1038/ncomms5157

Takehana, Y., Naruse, K., Hamaguchi, S., Sakaizumi, M., 2007. Evolution of ZZ/ZW and XX/XY sex-determination systems in the closely related medaka species, Oryzias hubbsi and O. dancena. Chromosoma 116, 463–470. https://doi.org/10.1007/s00412-007-0110-z

van Doorn, G.S., Kirkpatrick, M., 2007. Turnover of sex chromosomes induced by sexual conflict. Nature 449, 909–912. https://doi.org/10.1038/nature06178

Wada, H., Shimada, A., Fukamachi, S., Naruse, K., Shima, A., 1998. Sex-Linked Inheritance of the lf Locus in the Medaka Fish (Oryzias latipes). Zoolog. Sci. 15, 123–126. https://doi.org/10.2108/zsj.15.123

Wilson, C.A., High, S.K., McCluskey, B.M., Amores, A., Yan, Y., Titus, T.A., Anderson, J.L., Batzel, P., Carvan, M.J., III, Schartl, M., Postlethwait, J.H., 2014. Wild Sex in Zebrafish: Loss of the Natural Sex Determinant in Domesticated Strains. Genetics 198, 1291–1308. https://doi.org/10.1534/genetics.114.169284

Wright, A.E., Darolti, I., Bloch, N.I., Oostra, V., Sandkam, B., Buechel, S.D., Kolm, N., Breden, F., Vicoso, B., Mank, J.E., 2017. Convergent recombination suppression suggests role of sexual selection in guppy sex chromosome formation. Nat. Commun. 8, 14251. https://doi.org/10.1038/ncomms14251

Wright, A.E., Dean, R., Zimmer, F., Mank, J.E., 2016. How to make a sex chromosome. Nat. Commun. 7, 12087. https://doi.org/10.1038/ncomms12087

Yoshida, K., Makino, T., Yamaguchi, K., Shigenobu, S., Hasebe, M., Kawata, M., Kume, M., Mori, S., Peichel, C.L., Toyoda, A., Fujiyama, A., Kitano, J., 2014. Sex Chromosome Turnover Contributes to Genomic Divergence between Incipient Stickleback Species. PLOS Genet. 10, e1004223. https://doi.org/10.1371/journal.pgen.1004223

Zhou, Q., Bachtrog, D., 2012. Sex-Specific Adaptation Drives Early Sex Chromosome Evolution in Drosophila. Science 337, 341–345. https://doi.org/10.1126/science.1225385

