## Supplementary figures and images for "Diversity of sex chromosomes in Sulawesian medaka fishes"

### Figure S1

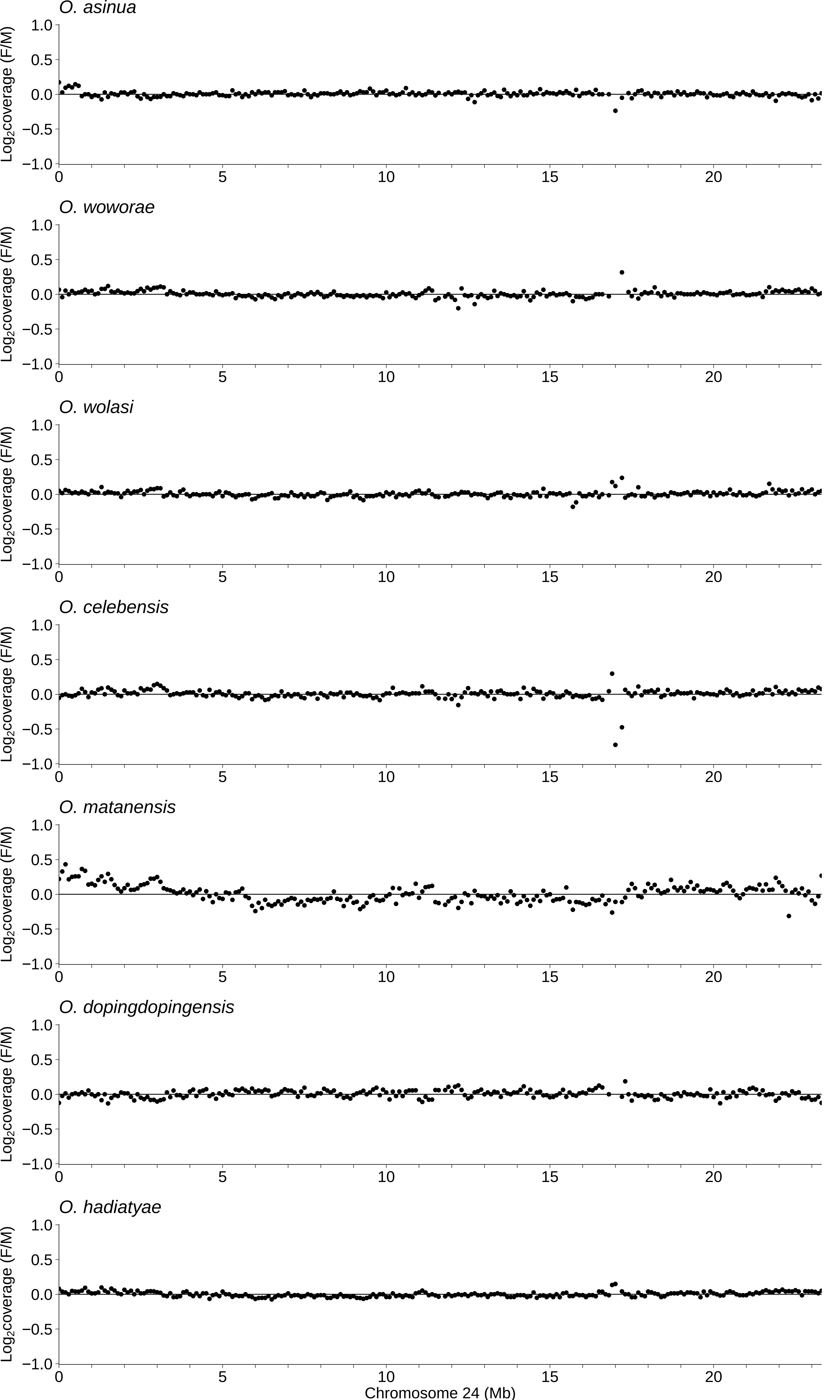
